# The Dishevelled C-terminus interacts with the centrosomal protein Kizuna to regulate microtubule organization during ciliogenesis

**DOI:** 10.1101/2025.11.12.688108

**Authors:** Maya Lines, Kenan Murray, Anthea Luo, Taewoo Yang, Yoo-Seok Hwang, Christopher J Westlake, Ira O. Daar, Jaeho Yoon

**Affiliations:** Cancer & Developmental Biology Laboratory, Center for Cancer Research, National Cancer Institute, National Institutes of Health, Frederick, MD 21702, USA; Laboratory of Cellular and Developmental Signaling, Center for Cancer Research, National Cancer Institute, National Institutes of Health, Frederick, MD 21702, USA

**Keywords:** Ciliogenesis, Dishevelled, microtubule meshwork, Kizuna

## Abstract

Microtubules (MTs) serve as dynamic scaffolds that support diverse cellular processes, including cell division, intracellular transport, and epithelial morphogenesis. In ciliated cells, MTs not only form the axonemal backbone of cilia but also work in tandem with actin to build an apical meshwork that anchors and orients basal bodies. However, the mechanisms regulating apical MT organization during ciliogenesis remain poorly understood. Here, we identify the centrosomal protein Kizuna (Kiz) as a critical regulator of apical MT architecture. Kiz interacts with the C-terminus of Dishevelled 2 (Dvl-C), inducing an open conformation and enabling the recruitment of Protein Kinase C delta (PKCδ) to stabilize the apical MT meshwork. This Dvl2–Kiz–PKCδ signaling axis is essential for ciliogenesis across multiple contexts, including the formation of primary cilia in the eye, multicilia in the mucociliary epidermis (MCE), and mono-motile cilia in the left-right organizer, the gastrocoel roof plate (GRP). These findings reveal a conserved molecular mechanism linking Dvl2 conformational dynamics to MT organization and highlight Kizuna as a key mediator coupling non-canonical Wnt signaling to ciliogenesis, left-right patterning, and the organization of the apical actin microtubule meshwork in ciliated cells of embryos.

## Introduction

Cilia are microtubule-based, antenna-like organelles that project from the surface of most eukaryotic cells. They are broadly classified into motile cilia, which include mono-motile and multiciliated forms that generate directional fluid flow in tissues such as the airway and reproductive tract(1, 2), and non-motile (primary) cilia, which are typically function as sensory organelles that detect chemical and mechanical cues to regulate intracellular signaling pathways(3). Extensive evidence indicates that cilia are indispensable for embryonic development and tissue homeostasis. Disruption of ciliogenesis or ciliary function results in a broad spectrum of human disorders collectively termed ciliopathies(4).

During ciliogenesis, apical cytoskeletal remodeling (including both actin and microtubule meshwork) is essential not only for proper basal body docking and axonemal extension but also for establishing basal body polarity and coordinating synchronized ciliary beating(5). Previously, we identified Developmentally regulated GTP-binding protein 1 (Drg1) as a novel Dishevelled (Dvl)-interacting protein essential for apical actin meshwork formation(6). Loss of Drg1 or disruption of its interaction with Dvl leads to shorter and fewer cilia, impaired basal body migration and docking, and defective rotational polarity in multiciliated cells. These defects stem from reduced apical actin meshwork formation and RhoA activity due to disrupted Dvl–Daam1–RhoA signaling.

Similar to the apical actin meshwork, microtubules also form an organized apical meshwork during ciliogenesis. The presence of a structured microtubule meshwork has been demonstrated using a GFP-tagged version of the ensconsin microtubule-binding domain (EMTB) in multiciliated cells (MCC) of Xenopus embryos(5). Treatment with the microtubule-depolymerizing drug nocodazole (Noc) disrupted the cytoplasmic microtubule meshwork resulting in defects of ciliary beating and polarity. It was proposed that the apical actin and microtubule meshworks play distinct and complementary roles during ciliogenesis(5). A cytoplasmic microtubule meshwork has also been observed in cells bearing primary cilia(7, 8). Microtubules nucleate around the basal body, which corresponds to the centriole, serving as the primary microtubule-organizing center (MTOC). Despite the presence of an apical microtubule meshwork, its specific functions and the molecular mechanisms governing its organization remain poorly understood.

Dishevelled (Dvl), a central scaffold protein in the Wnt signaling pathway, has been shown to play an essential role in ciliogenesis. It was revealed that depletion of Dvl leads to profound defects in cilia formation, characterized by aberrant apical docking and disrupted polarity of basal bodies. Interestingly, the C-terminal region of Dvl2 (Dvl2-C), which lacks the conserved DIX, PDZ, and DEP domains, is sufficient for targeting Dvl to the basal bodies, suggesting that this region contains a distinct functional module critical for ciliogenesis(9). This Dvl2-C region is highly conserved across Dvl isoforms and among species from humans to fish. Our previous work demonstrated that Wnt–PCP components, including Ror2 and Shroom3, interact with Dvl2-C to modulate actomyosin contractility during neural plate folding, further supporting the idea that the Dvl2 C-terminus serves as a unique regulatory module with functions that may extend to ciliogenesis(10).

Here, we identified Kizuna (also known as polo-like kinase 1 substrate) as a novel Dvl2-C–binding partner that plays a critical role in organizing the apical microtubule meshwork during ciliogenesis. Functional analyses revealed that loss of Kizuna caused severe defects in multicilia formation, disrupted left–right patterning due to abnormal mono-motile cilia in the gastrocoel roof plate (GRP), and retinal pigment epithelium (RPE) malformation associated with impaired primary cilia. Furthermore, we demonstrate that Kizuna interacts with Dvl2-C and recruits PKC-δ, whose kinase activity enhances the stability of pericentriolar material (PCM), thereby promoting apical microtubule meshwork assembly and proper cilia formation.

## Results

### Kizuna, a Novel Dvl2-C Binding Partner, Is Expressed in Ciliated Cells

To elucidate the molecular mechanism by which Dvl2-C contributes to ciliogenesis, we conducted an immunoprecipitation–mass spectrometry (IP/MS) screen using HA–Dvl2-C that was overexpressed in Xenopus embryos to identify its interacting partners (Fig 1a). Among the candidates, Kizuna was identified as a promising Dvl2-C–binding partner. Kizuna is a well-characterized component of the pericentriolar material (PCM) that localizes to the PCM during mitosis in HeLa cells, where it functions to stabilize microtubule organization(7). Subsequent studies have shown that Kizuna also localizes to the basal body in primary cilia(11), suggesting an additional role beyond cell division. Moreover, bulk RNA-seq analysis revealed that Kizuna is expressed in the gastrocoel roof plate (GRP)(12), a mono-motile ciliated structure often associated with establishment of proper left-right organ patterning. Together, these findings highlight Kizuna as a compelling candidate for a novel regulator of ciliogenesis acting through its interaction with Dvl2-C.

**Figure 1.**
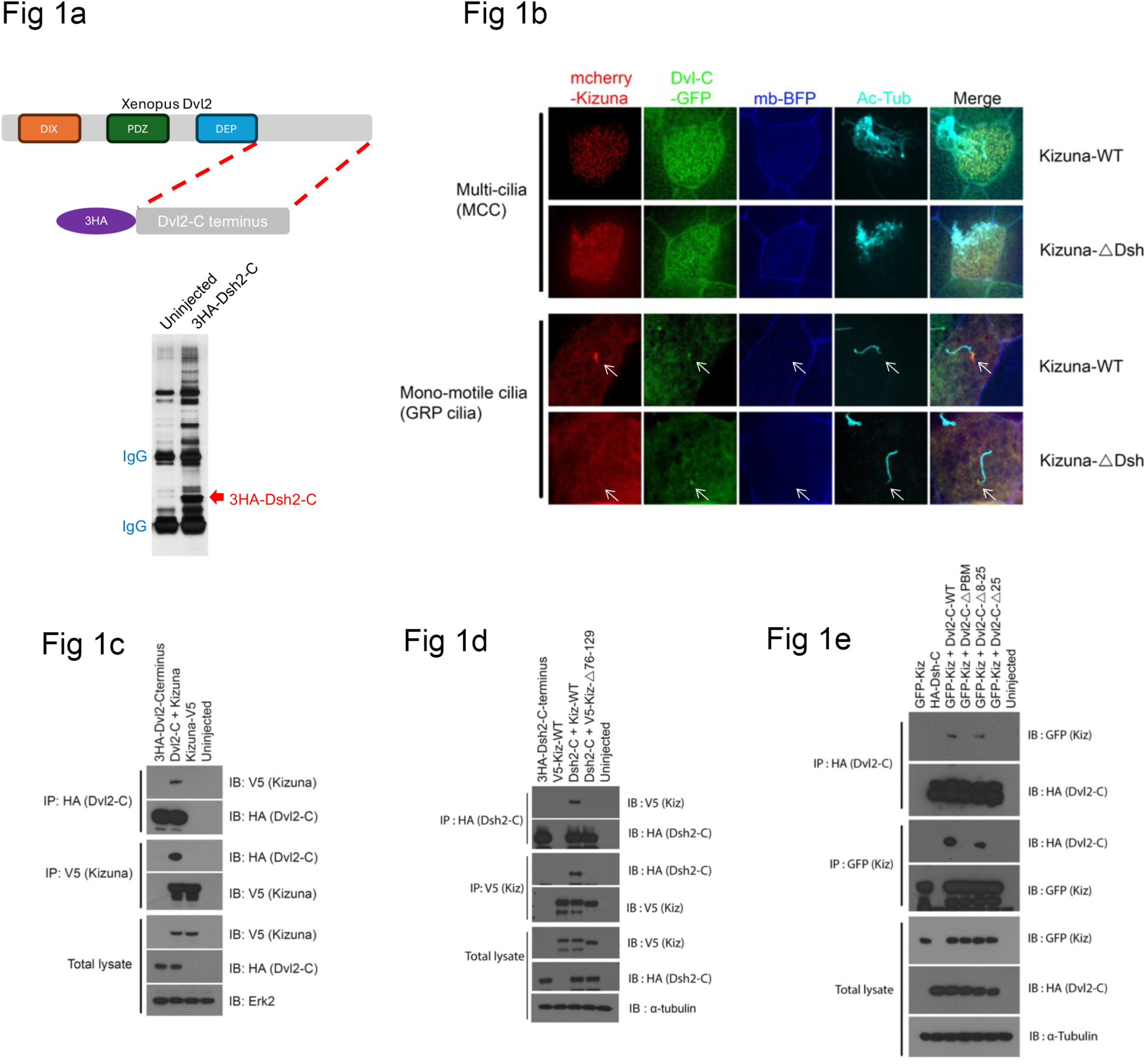
Kizuna interacts with Dvl2 through its N-terminal domain and co-localizes at basal bodies. (a) Schematic representation of Dvl2 conserved domains and the HA-tagged Dvl-C construct used for pull-down assays in Xenopus embryos. See also Supplementary Table 1. (b) Co-localization of Kizuna and Dvl-C in multiciliated cells (MCCs) and mono-motile ciliated cells of the gastrocoel roof plate (GRP). (c) Co-immunoprecipitation (Co-IP) of HA–Dvl-C (200 pg) with V5–Kizuna (1 ng) expressed in Xenopus embryos. (d) Co-IP analysis of HA–Dvl-C with a Kizuna deletion mutant lacking amino acids 76–129. (e) Co-IP of full-length Kizuna with serial Dvl-C deletion mutants. See also Figure S1d.

Expression analysis revealed Kizuna enrichment in developing multiciliated cells (MCCs), as shown by reanalysis of single-cell RNA-seq data from the Kedar Nath Natarajan laboratory (Fig. S1a). Immunostaining confirmed that Kizuna localizes around basal bodies in MCCs and mono-motile ciliated cells in GRP, a pattern similar to that of Dvl2-C (Fig 1b). In situ hybridization further showed that Kizuna is expressed in the GRP, the left–right organizer containing mono-motile ciliated cells (Fig S1b).

We next validated the Dvl2-C/Kizuna interaction by co-immunoprecipitation (Co-IP) using Kizuna-overexpressing Xenopus embryos (Fig. 1c). Deletion analysis revealed that the N-terminal region of Kizuna, specifically residues 76–129, is required for Dvl2-C binding (Fig 1d and Fig S1c). Conversely, Co-IP assays with Dvl2-C deletion mutants demonstrated that the C-terminal six amino acids, corresponding to the PDZ-binding motif (PBM), are essential for interaction with Kizuna (Fig 1e and Fig S1d). These results indicate that the C-terminal tail of Dvl2 interacts with the N-terminal domain of Kizuna. To determine whether the Dvl2-C/Kizuna interaction is required for Kizuna’s basal body localization, we examined Kizuna deletion mutants lacking amino acids 76–129 (Kizuna-ΔDvl), which abolish Dvl2-C binding. Unlike wild-type Kizuna (Kizuna-WT), the Kizuna-ΔDvl mutant failed to localize to basal bodies (Fig 1b).

Since Kizuna is phosphorylated by PLK1 during mitosis, we assessed whether this modification influences its interaction with Dvl2-C or localization. Co-IP analysis showed that both non-phosphorylatable (T369A) and phosphomimetic (T369E) mutants interacted with Dvl2-C at levels comparable to wild type (Fig S1e). Consistently, both mutants exhibited similar subcellular localization (Fig S1f), indicating that PLK1-mediated phosphorylation does not affect the Dvl2-C/Kizuna interaction or basal body targeting. These results suggest that Kizuna associates with basal bodies through its interaction with Dvl2-C and that this interaction may be important for proper ciliogenesis.

### Loss of Kizuna leads to ciliogenesis defects

Having established that Kizuna interacts with Dvl-C and co-localizes near the basal body in ciliated cells, we next examined whether cilia formation is affected by Kizuna depletion. To define the functional role of Kizuna, we designed a specific morpholino (MO) targeting Kizuna and confirmed its efficiency by blocking the translation of exogenous Kizuna protein (Fig. S2a). To knock down Kizuna in multiciliated cells (MCCs), the MO was injected into the V1.1 blastomere at the 8-cell stage. Immunostaining analysis revealed that cilia, visualized with acetylated tubulin, were significantly reduced in the MCCs of Kizuna morphant embryos (Fig. 2a and Fig. S2c), resembling the ciliogenesis defects observed in Dvl morphants(9). Similarly, CRISPR/Cas12-mediated genetic deletion of Kizuna produced comparable ciliogenesis defects in F0 crispant embryos, confirming that the phenotype was specifically due to loss of Kizuna function (Fig. S2b). The ciliary defects in Kizuna morphants were rescued by re-expression of MO-resistant wild-type (WT) Kizuna but not by the Dvl-C–binding–deficient mutant Kizuna-ΔDvl. Live imaging further demonstrated that, while control embryos displayed robust and synchronized multiciliary beating, Kizuna knockdown severely impaired both cilia formation and coordinated motility (Supplementary Movie1). These findings indicate that Kizuna is essential for multicilia formation and that its interaction with Dvl-C is critical for proper multi-ciliogenesis.

**Figure 2.**
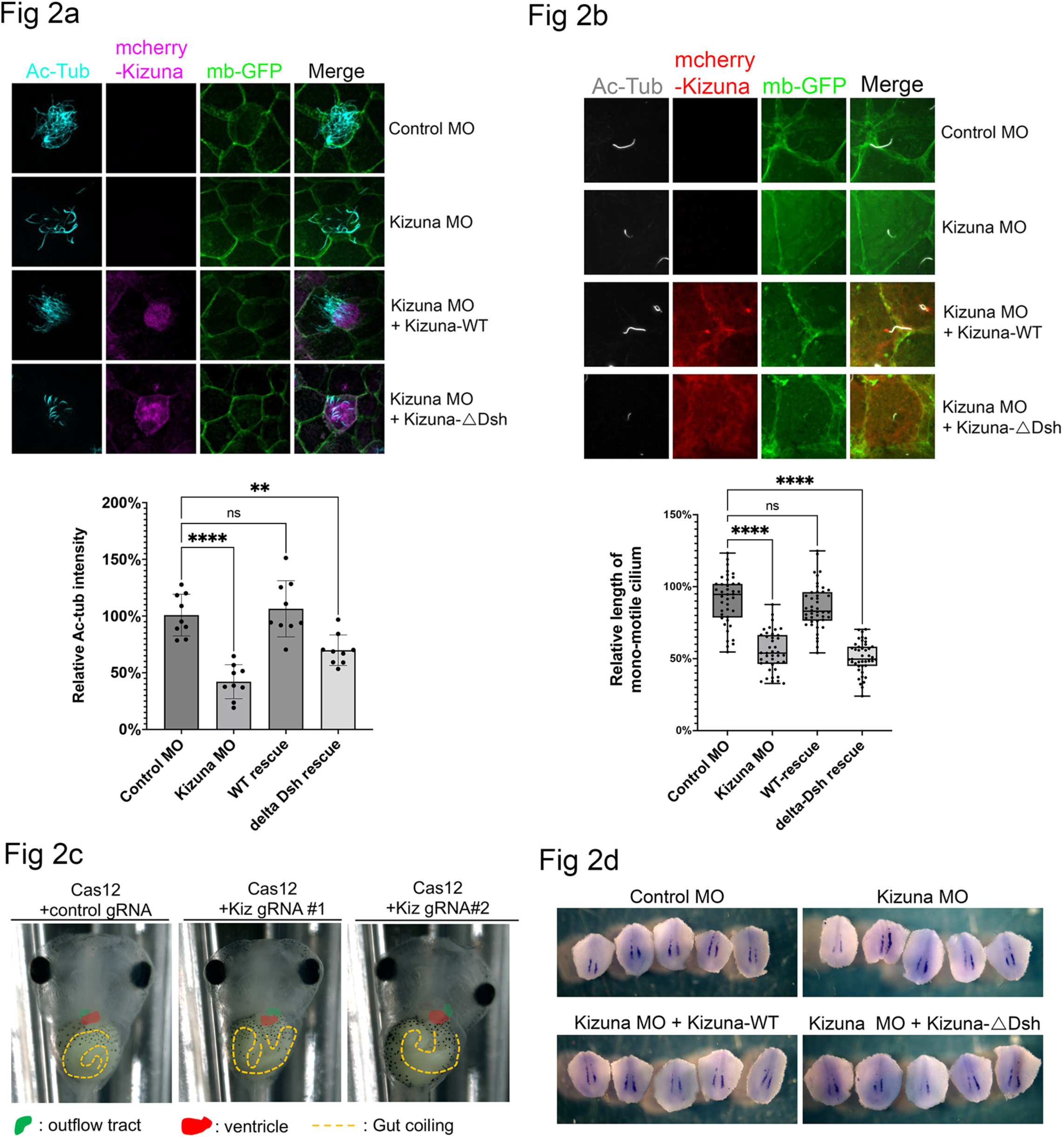
Kizuna is essential for multiciliogenesis, mono-motile cilium formation. (a) Immunostaining of multiciliated cells (MCCs) in Xenopus embryos at stage 30 following Kizuna morpholino (MO) injection. The box plot represents minimum to maximum values with all data points shown. See also Fig. S2c for low-magnification views. (b) Immunostaining of mono-motile ciliated cells in dissected gastrocoel roof plate (GRP) tissues from Kizuna morphants at stage 20. The box plot represents minimum to maximum values with all data points shown. See also Fig. S2d and e for low-magnification views. (c) Representative phenotypes of Kizuna CRISPR knockout embryos showing inverted heart looping and reversed gut coiling, indicative of left–right asymmetry defects. (d) *In situ* hybridization with dand5 (also known as Coco) probe in Kizuna morphants reveals disrupted asymmetric gene expression at the GRP.

The requirement for Kizuna function in multicilia begs the question of whether it plays a role in mono-motile cilia that are found in the GRP. Therefore, we specifically targeted the GRP by injecting Kizuna morpholinos into the dorsal marginal zone of two blastomeres at the 4-cell stage. Knockdown of Kizuna resulted in a ∼50% reduction in mono-motile cilium length (Fig. 2b, Fig S2d-e, and Supplementary Movie2). This defect was rescued by re-expression of wild-type Kizuna but not by the Dvl-C–binding–deficient mutant Kizuna-ΔDvl, indicating that the Dvl-C/Kizuna interaction is required for proper cilium formation. Mono-motile cilia in the gastrocoel roof plate (GRP) are associated with the establishment left–right (LR) body asymmetry(2, 13). In Xenopus embryos, LR symmetry breaking correlates with a leftward fluid flow generated by the coordinated rotation of mono-motile cilia within the GRP, which functions as the left–right organizer (LRO). Defective formation or function of these cilia disrupts this directional flow and is often associated with heterotaxy (HTX), characterized by abnormal positioning or morphology of internal organs along the LR axis(14, 15). Consistent with these ciliary defects, Kizuna morphants exhibited heterotaxy phenotypes, including reversed heart and gut looping (Fig. 2c). At the molecular level, the leftward flow triggers asymmetric Ca²⁺ signaling, leading to unilateral activation of the Nodal pathway(16). The Nodal antagonist dand5 is expressed at the marginal zone of the LRO, and its repression on the left side enables left-sided activation of Nodal–Pitx2 signaling, which ultimately determines organ situs during organogenesis(15, 17–20). Consistent with the mono-motile cilia defects observed in Kizuna morphants, loss of Kizuna disrupted the asymmetric expression of dand5 (Fig. 2d) and resulted in inverted heart and gut looping defects. These findings demonstrate that Kizuna is essential for both multiciliogenesis and mono-motile cilium formation and plays a critical role in establishing left–right asymmetry during embryogenesis.

### Kizuna regulates apical microtubule meshwork formation during ciliogenesis

Since Kizuna has been shown to localize to the pericentriolar material (PCM) and stabilize microtubule organization during mitosis in cell culture(8), we next tested whether it also contributes to the formation of the apical microtubule meshwork in ciliated cells. Microtubules play diverse and highly coordinated roles in ciliated cells. They form an apical meshwork beneath the cilia and interact with actin filaments to regulate apical surface expansion following intercalation of multiciliated cells (MCC)(5, 21). The apical microtubule meshwork also helps maintain apical surface architecture and provides mechanical resistance against deformation during ciliary beating(22). Moreover, it contributes to the precise orientation of basal bodies, which is essential for coordinated and synchronized ciliary motion(5, 6). As shown in Fig. 3a, an apical microtubule meshwork forms following MCC intercalation during Xenopus skin development. This apical MT meshwork formation closely resembles the apical actin meshwork and is positioned to support apical surface expansion and organization. To assess the role of the apical microtubule (MT) meshwork in ciliogenesis, embryos were treated with the MT-depolymerizing drug nocodazole after multicilia formation (stages 24–30). Nocodazole treatment markedly disrupted the apical MT meshwork and led to defective cilia formation, indicating that continuous microtubule polymerization is required (Fig S3a).

**Figure 3.**
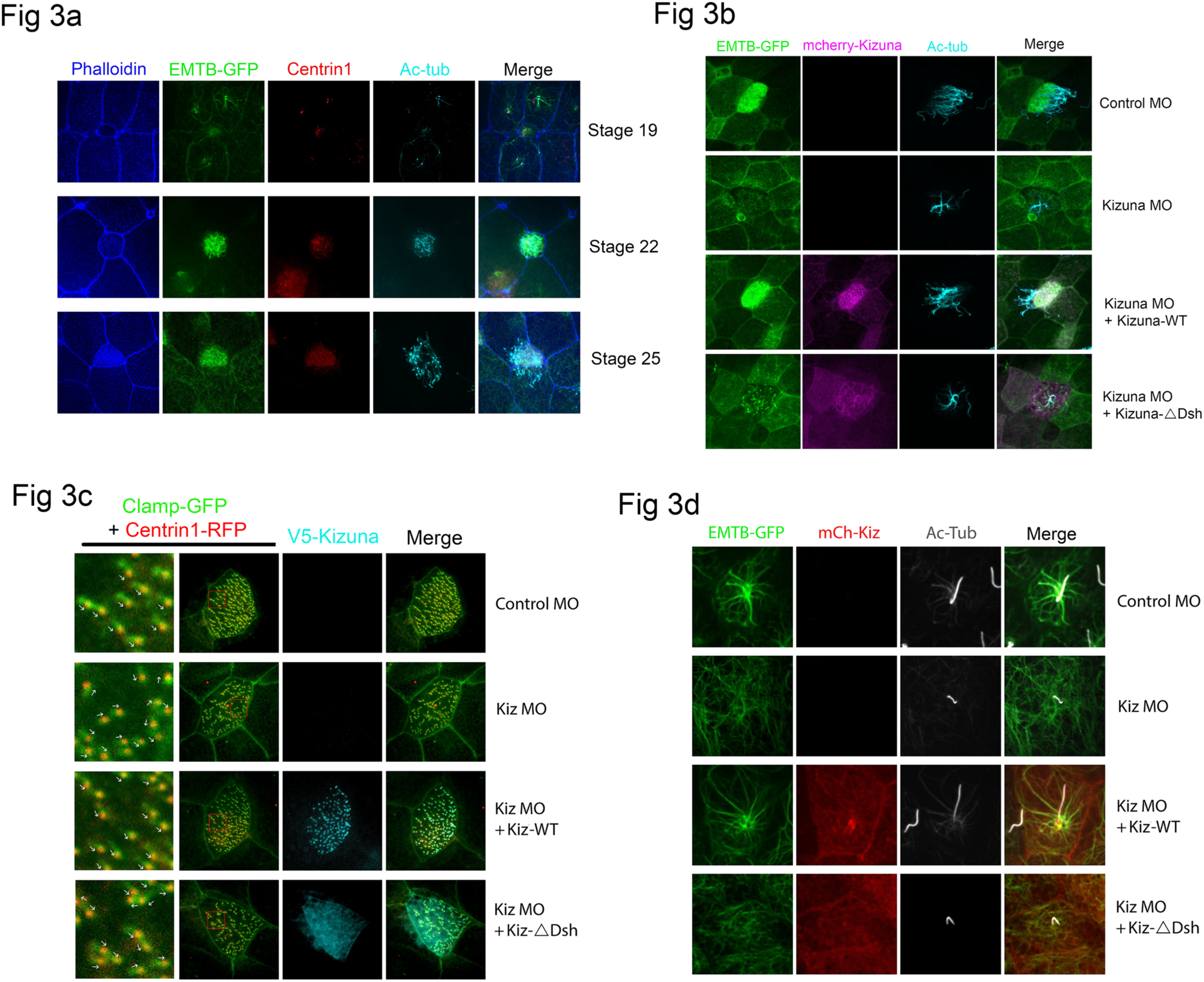
Kizuna regulates microtubule meshwork organization. (a) Immunostaining of the apical microtubule (MT) meshwork in developing multiciliated cells (MCCs). (b) Immunostaining of the apical MT meshwork in Kizuna morphants at stage 30. See also Fig. S3b for low-magnification views. (c) Analysis of cilia polarity using overexpressed Clamp-GFP and Centrin1-RFP in MCCs of Xenopus embryos at stage 30. (d) Immunostaining of microtubule nucleation in mono-motile ciliated cells within the gastrocoel roof plate (GRP) at stage 20. See also Fig. S3d for low-magnification views.

Knockdown of Kizuna markedly disrupted the apical microtubule (MT) meshwork in multiciliated cells (MCCs), resulting in shorter and fewer cilia. These defects were rescued by re-expression of wild-type Kizuna but not by the Dvl-C–binding–deficient mutant Kizuna-ΔDvl (Fig 3b and S3b). In contrast, loss of Kizuna did not affect actin meshwork formation or basal body apical docking, suggesting that Kizuna specifically regulates MT organization (Fig S3c). Because microtubules play a key role in establishing basal body polarity, we next examined whether Kizuna depletion influences this process. Disruption of MT organization led to randomized cilia orientation. Indeed, Kizuna knockdown caused a loss of planar polarity in MCCs, which was rescued by wild-type Kizuna but not by Kizuna-ΔDvl (Fig 3c). We further analyzed the MT meshwork in mono-motile ciliated cells of the gastrocoel roof plate (GRP) and observed distinct MT nucleation around the basal body, which was dramatically reduced upon Kizuna depletion. Reintroduction of wild-type Kizuna restored MT nucleation, whereas Kizuna-ΔDvl failed to do so (Fig. 3d and S3d). Together, these findings demonstrate that the Dvl-C/Kizuna interaction is essential for maintaining microtubule architecture during ciliogenesis.

### The Dvl-C/Kizuna complex recruits PKCδ to promote microtubule meshwork stabilization

PKCδ has been implicated in the non-canonical Wnt signaling pathway and is known to interact with Dishevelled (Dvl)(23). In addition, PKCδ associates with microtubule-organizing center (MTOC) components and regulates microtubule nucleation and spindle organization during mitosis(8). Its kinase activity has also been shown to maintain the integrity of the pericentriolar material (PCM), a key microtubule-nucleating structure. To investigate how PKCδ contributes to apical MT organization, we analyzed its subcellular localization. PKCδ was enriched at the basal body in MCCs, and a similar localization was observed in mono-motile cilia within the GRP (Fig 4a-b). Interestingly, Kizuna knockdown suppressed PKCδ basal body localization, whereas re-expression of wild-type Kizuna restored it, but not the Kizuna-ΔDvl mutant (Fig 4a-b). Next, we tested whether Kizuna influences the Dvl/PKCδ interaction. Overexpression of Kizuna enhanced Dvl–PKCδ binding, whereas the Kizuna-ΔDvl failed to do so (Fig 4c). Our previous study demonstrated that the PDZ-binding motif (PBM), which mediates Kizuna binding, is critical for Dvl’s conformational switch between open and closed states. This raises the possibility that Kizuna binding to Dvl-PBM promotes the open conformation of Dvl, exposing the DEP domain for PKCδ interaction. To evaluate this idea, we expressed the PBM-deletion or PDZ domain deletion Dvl mutant that adopts a constitutively open conformation and compared its interaction with PKCδ to that of wild-type Dvl. Remarkably, constitutively open conformation Dvl (PBM deletion or PDZ domain deletion mutants) increased Dvl–PKCδ binding even without Kizuna overexpression (Fig 4d-e).

**Figure 4.**
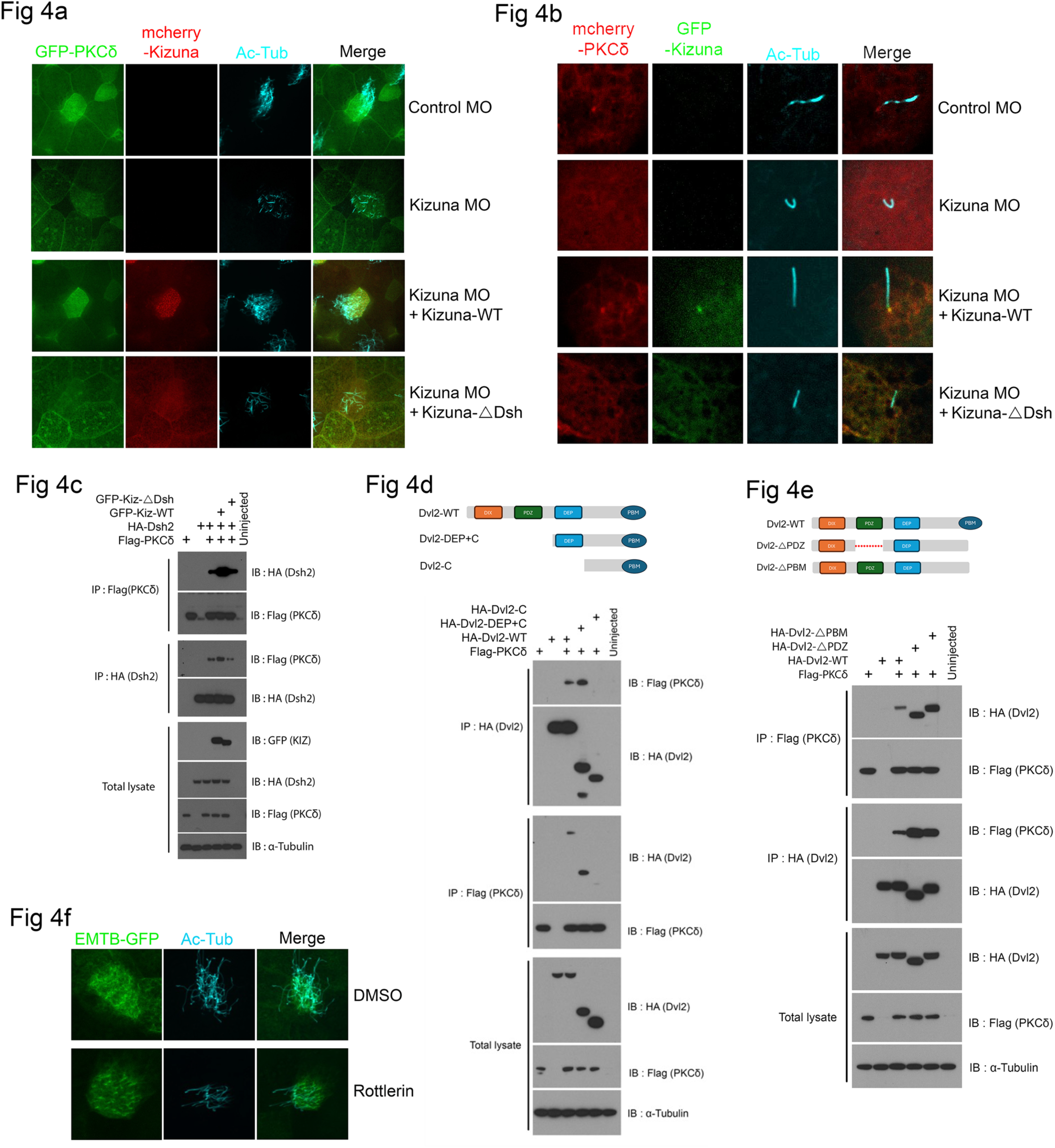
PKCδ acts downstream of the Dvl2/Kizuna complex. (a) Subcellular localization of PKCδ in MCCs at stage 30. See also Fig. S4a for low-magnification views. (b) Subcellular localization of PKCδ in mono-motile ciliated cells at stage 20. (c) Co-immunoprecipitation (Co-IP) of PKCδ with WT-Dvl2 in the presence of WT-Kizuna or Kizuna-ΔDvl mutant. (d) Co-IP of PKCδ with serial deletion constructs of Dvl2 as shown in the schematic representation. (e) Co-IP of PKCδ with Dvl2 deletion mutants lacking the PDZ domain or PBM motif. (f) Immunostaining of the apical MT meshwork following rottlerin (10 μM) treatment from stage 22 to 27.

Given that Kizuna contributes to PKCδ recruitment to Dvl and is required for the apical microtubule meshwork in multiciliated cells (MCCs) and to microtubule nucleation in mono-motile cilia of the gastrocoel roof plate (GRP), we next investigated whether PKCδ is involved in Kizuna-mediated signaling. Since knockdown of PKCδ by morpholino oligos or overexpression of dominant-negative PKCδ caused severe developmental defects at early stages of Xenopus embryos, we instead used Rottlerin to inhibit PKCδ kinase activity from stage 22 to 27. As shown in Fig. 4f, Rotterlin treatment dramatically suppressed apical microtubule (MT) meshwork formation and multicilia development in MCCs. These findings strongly suggest that PKCδ activity is essential for apical MT meshwork formation during ciliogenesis. Collectively, these results suggest that Kizuna associates with the Dvl-PBM region, inducing an open Dvl conformation that allows PKCδ to access the DEP domain. PKCδ activity, in turn, regulates microtubule meshwork formation during ciliogenesis.

### Kizuna depletion disrupts primary cilium formation in RPE1 cells, contributing to retinal degeneration in retinitis pigmentosa

In human patients, mutations in the Kizuna gene are associated with retinitis pigmentosa, a degenerative retinal disease linked to impaired retinal pigment epithelium (RPE) differentiation, a process tightly regulated by primary cilia(11). To explore the clinical relevance of Kizuna deficiency, we injected Kizuna-specific morpholino (MO) unilaterally into the D1.1.1 blastomere, which contributes to more than 50% of the retinal population at the 32-cell stage. Consistent with human patient phenotypes, Kizuna knockdown resulted in a marked reduction of eye pigmentation in developing Xenopus embryos (Fig. 5a). Cross-sectional immunostaining further revealed that Kizuna depletion dramatically decreased RPE65 expression, a marker of differentiated RPE cells (Fig 5b). Co-staining with acetylated tubulin and RPE65 showed that loss of Kizuna severely reduced acetylated tubulin staining in the RPE layer, indicating that ciliogenesis defects may underlie the observed RPE differentiation failure (Fig 5c). To rule out the possibility that reduced pigmentation arose from apoptosis in the developing eye, we examined cleaved caspase-3 levels. No significant increase in cleaved caspase-3 was detected in the RPE layer, suggesting that cell death is not the primary cause of the phenotype (Fig S5). Kizuna function was further validated in human RPE1 cell culture, where Kizuna knockdown by siRNA led to approximately 50% reduction in primary cilium length (Fig 5d).

**Figure 5.**
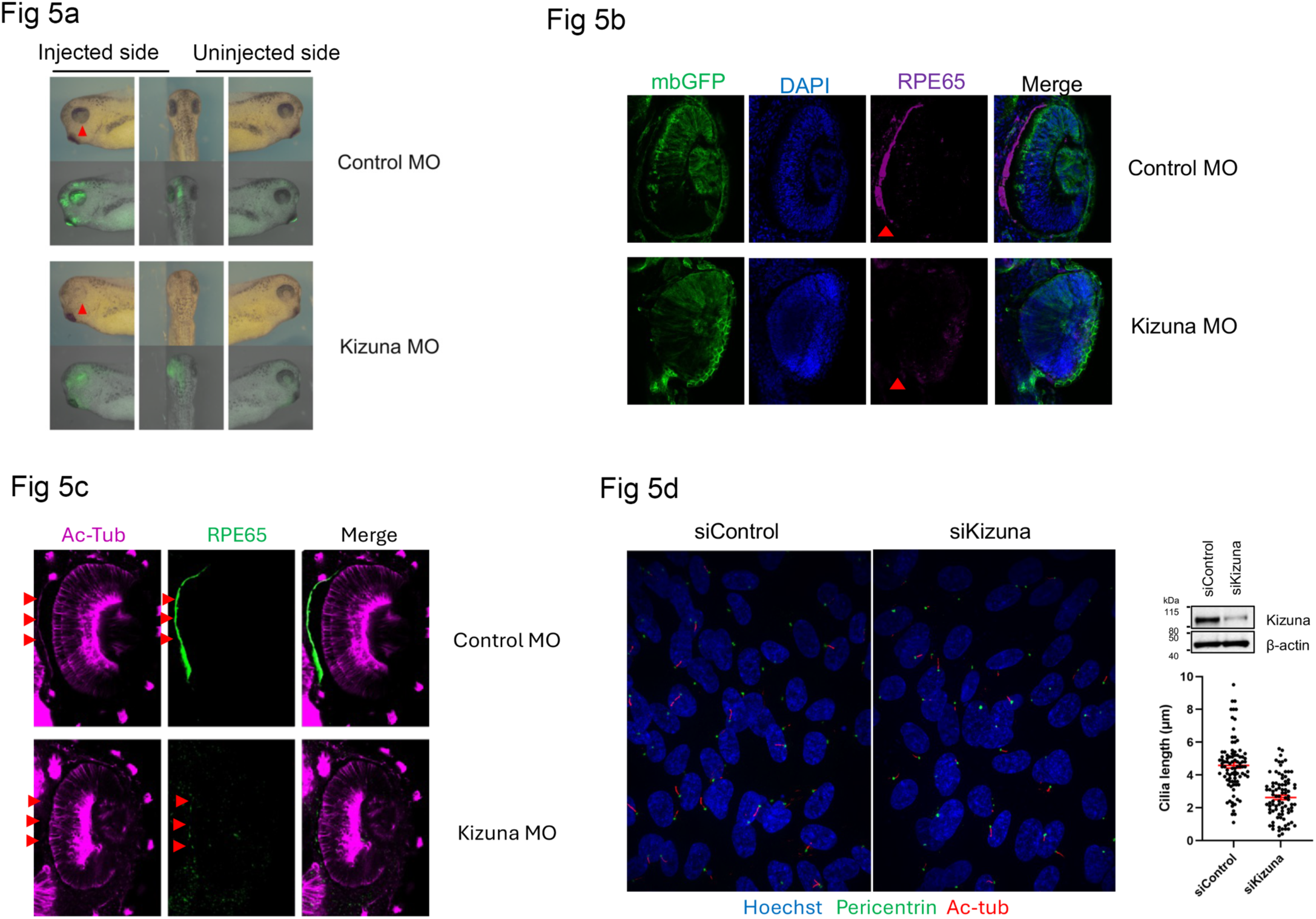
Kizuna regulates retinal pigment epithelium (RPE) differentiation and ciliogenesis. (a) Representative phenotypes of Xenopus embryos at stage 30 following unilateral Kizuna morpholino (MO) injection into the D1.1 blastomere at the 8-cell stage. (b) Immunostaining of the RPE layer marker RPE65 in eyes of Kizuna MO-injected embryos at stage 35. (c) Co-staining of RPE65 and acetylated tubulin in eyes of Kizuna MO-injected embryos at stage 35. (d) Immunostaining of primary cilia in Kizuna knockdown (siRNA) RPE1 cells. Immunoblot shows endogenous Kizuna expression. The dot plot displays all data points.

Together, these findings demonstrate that Kizuna is also essential for primary cilium formation, and its loss disrupts RPE differentiation, providing a mechanistic explanation for the retinitis pigmentosa in patients with Kizuna mutations.

## Discussion

Ciliogenesis requires precise coordination between cytoskeletal networks and signaling pathways. While the apical actin meshwork has been extensively characterized by multiple groups, the organization and function of the apical microtubule (MT) network remain understudied. Previous reports using nocodazole disruption have suggested that MTs contribute to multiciliated cell (MCC) radial intercalation and cilia polarity; however, the mechanisms governing apical MT meshwork assembly are still poorly understood. In this work, we identified the centrosomal protein Kizuna (Kiz) as a novel binding partner of Dishevelled 2 (Dvl2), a central scaffold in Wnt signaling using Xenopus laevis embryos. Immunoprecipitation–mass spectrometry revealed that Kiz interacts with the C-terminal PDZ-binding motif (PBM) of Dvl2 via its N-terminal domain. Kiz localizes to basal bodies in both MCCs and mono-motile cilia of the gastrocoel roof plate (GRP), and this localization requires its interaction with Dvl2. Loss-of-function analyses demonstrated that Kiz is essential for the formation of both multicilia and mono-motile cilia. Our findings suggest that the mechanisms governing spindle assembly during mitosis and apical MT meshwork formation during ciliogenesis may share conserved organizational principles.

We further show that Protein Kinase C delta (PKCδ), a kinase activity implicated in non-canonical Wnt signaling and PCM integrity, joins and acts in concert with the Dvl2–Kiz complex. Inhibition of PKCδ activity disrupted apical MT meshwork and cilia formation, demonstrating its essential role in ciliogenesis. Kiz knockdown abolished PKCδ basal body localization, while wild-type Kiz rescued it, but a Dvl-binding mutant did not. Mechanistically, Kiz binding to the PBM motif of Dvl-caboxyl terminus induces an open Dvl conformation that exposes the DEP domain, facilitating PKCδ binding and localization at the basal body. In addition, the clinical relevance of Kizuna was validated in Xenopus and human RPE1 cells. Knockdown of Kizuna in Xenopus reduced retinal pigment epithelium (RPE) differentiation and primary cilia formation, phenocopying human KIZ mutations associated with retinitis pigmentosa.

Collectively, these findings reveal a conserved Dvl2–Kiz–PKCδ signaling axis that regulates apical microtubule organization and ciliogenesis. This pathway links non-canonical Wnt signaling to cytoskeletal remodeling, providing new insights into planar cell polarity–mediated developmental processes and ciliopathy-associated diseases.

## Materials and Methods

### Xenopus laevis

Wild-type Xenopus laevis were purchased from Nasco (USA). All experimental procedures were reviewed and approved by the Animal Care and Use Committee of the National Cancer Institute-Frederick (ASP #18-433) and performed in compliance with AAALAC standards.

### Plasmids and mRNA synthesis

The full-length Xenopus Kizuna cDNA clone was obtained from Horizon (GenBank ID: BC129527). Deletion constructs and point mutants (Kizuna-T369A and Kizuna-T369E) were generated in the pCS107 vector using the QuikChange II Site-Directed Mutagenesis Kit. Capped sense RNAs were synthesized with the mMESSAGE mMACHINE SP6 Kit (Thermo Fisher) following linearization with ASP718.

### Embryonic injections

Xenopus embryos were obtained using standard methods (Moody, 2000). Morpholino oligonucleotides (MOs) and mRNAs were microinjected into specific blastomeres at defined developmental stages: into the animal pole region at the one-cell stage for CRISPR/Cas12 knockout, into the dorsal marginal zones at the four-cell stage to target gastrocoel roof plates (GRP), into the V1 blastomere at the eight-cell stage to target epidermal tissue, or into the D1.1 blastomere to target the eye. All experimental procedures were approved by the Animal Care and Use Committee of the National Cancer Institute-Frederick (ASP #18-433) and were conducted in compliance with AAALAC guidelines.

### Whole-mount in situ hybridization (WISH)

Dissected GRP were collected at stage 20 for hybridization with the anti-Kizuna or dand5 probes. Embryos were injected with membrane-GFP and various mRNAs or MOs to distinguish the injected side of embryos. The embryos were then processed for whole-mount in situ hybridization using standard methods (Harland, 1991).

### Co-immunoprecipitation and Western blot analysis

Embryos were lysed in lysis buffer (50 mM Tris-HCl [pH 7.4], 150 mM NaCl, 1% NP-40, 0.5 mM phenylmethyl sulphonyl fluoride (PMSF, ThermoFisher), protease inhibitor cocktail (Roche), and phosphatase inhibitors (100mM Sodium Vanadate and 10mM b-glycerophosphate). The cell lysates were cleared by centrifugation at 13,000 g for 10 min at 4 °C. IPs were performed at 4 °C for 8 h with the following agarose beads: Anti-HA-agarose (Sigma-Aldrich), Anti-Flag-agarose (Sigma-Aldrich), Anti-myc-agarose (Sigma-Aldrich), GFP-Trap affinity resin (Chromotek), and GFP-V5 affinity resin (Chromotek). Western blot analysis was performed using anti-Flag-HRP-conjugated (1:5,000, Sigma-Aldrich), anti-HA-HRP-conjugated (1:5,000, Sigma-Aldrich), anti-myc-HRP-conjugated (1:5,000, Sigma-Aldrich), anti-GFP-HRP-conjugated (1:5,000, Rockland Immunochemicals), anti-V5-HRP-conjugated (1:5,000, ThermoFisher), anti-ERK2 (1:1000, Santa Cruz Biotechnology), anti-a-tubulin (1:1000, Abcam) and mouse anti-alpha-tubulin-HRP-conjugated (Proteintech). Secondary antibodies used were goat anti-rabbit-HRP-conjugated (Cell Signaling Technology) and mouse anti-goat-HRP-conjugated (Santa Cruz Biotechnology).

### Immunofluorescence and live imaging

Immunofluorescence analyses were performed on dissected GRPs at stage 20 or on tadpoles as previously described. Briefly, explants were fixed in MEMFA (4% formaldehyde in 1× MEM salts) overnight at 4 °C or in 2% trichloroacetic acid (TCA) for 30 min at room temperature, followed by dehydration in 100% methanol. Samples were blocked in 10% goat serum in 1× PBS (filtered) and incubated with the following primary antibodies: mouse anti–acetylated tubulin (1:1000, T7451, Sigma), mouse anti–γ-tubulin (1:500, ab113717, Abcam), mouse anti–HA–Alexa Fluor 488/555/647 (1:500, 2-2.2.14, Thermo Fisher), mouse anti–V5–Alexa Fluor 488/555/647 (1:500, 2F11F7, Thermo Fisher), and rabbit anti–GFP–Alexa Fluor 488 (1:500, A-21311, Thermo Fisher). Secondary antibodies included Alexa Fluor 488 or 594–conjugated goat anti–mouse or anti–rabbit IgG (1:500, Invitrogen). F-actin was visualized with Alexa Fluor 488–phalloidin (A12379, Thermo Fisher). Imaging was performed using a Marianas spinning-disk confocal microscope (Intelligent Imaging Innovations) equipped with 40×/1.4 NA or 63×/1.3 NA objectives. Images were acquired and processed using SlideBook (Intelligent Imaging Innovations) or ZEN (Zeiss) software. Time-lapse imaging of fluorescently tagged proteins was conducted as previously described and detailed in the figure legends. For cilia-beating analyses, stage 30 embryos were anesthetized in 150 mg/L tricaine (MS-222; Sigma, A-5040), and embryos with optimal fluorescence were imaged at room temperature using the Marianas system with a 63×/1.3 NA objective.

### CRISPR/Cas12a genome editing

Guide RNA (gRNA) were designed using the CRISPRscan web tool (https://www.crisprscan.org) to identify suitable Cas12a target sites in the Xenopus laevis genome. A target sequence within exon 2 (TTTCCAGCACAGCTCATCTGATGCACA) was selected. The gRNA and Alt-R LbCas12a (Cpf1) protein were purchased from Integrated DNA Technologies (IDT). A Cas12a ribonucleoprotein (RNP) complex containing 10 µM gRNA and 10 µM Cas12a was prepared, and 5 nL of the mixture was injected into the animal pole region of one-cell-stage embryos. Injected embryos were cultured at 20 °C until the desired developmental stages. To confirm genome editing, F0 embryos were imaged and subsequently analyzed by direct sequencing of PCR amplicons spanning the targeted region (F: 5’-GGTAAGTTCAGAGGCATGCC-3’, R: 5’-CTGCCCCTAAAAGTACCCCT-3’).

### Statistical analysis

Sample sizes are indicated in the corresponding figure legends. No statistical method was used to predetermine sample size. Dead cells and embryos were excluded from all analyses. Embryos with mistargeted injections were also excluded; targeting accuracy was confirmed by co-injection of lineage tracers (membrane-GFP or membrane-RFP RNAs). All experiments were performed in a blinded manner, and the order of testing was randomized. Pilot experiments were conducted to determine optimal RNA and morpholino dosages, followed by full-scale experiments performed independently at least three times. Quantifications were performed using ImageJ. Data normality was assessed using the Kolmogorov–Smirnov, D’Agostino–Pearson omnibus, and Shapiro–Wilk tests in Prism 10; datasets were considered normally distributed only if all three tests indicated normality. For normally distributed data, comparisons were made using a two-tailed Student’s t-test (unequal variances) or one-way ANOVA with Dunnett’s multiple comparisons post-test in Prism 10. Non-normally distributed data were analyzed using the Mann–Whitney test or nonparametric ANOVA (Kruskal–Wallis test with Dunn’s multiple comparisons post-test) in Prism 9. Cross-comparisons were performed only when the overall P value from ANOVA was < 0.05. Error bars represent the standard error of the mean (SEM).

## Acknowledgements

We thank S. Lockett for technical support for microscopy, and all members of the Cancer and Developmental Biology Laboratory for discussion and comments. We are grateful to the K. Natarajan laboratory for single-cell-RNA-seq analysis. This research was supported by the National Cancer Institute, National Institutes of Health, Intramural Research Program, and was funded in whole or in part with federal funds from the National Cancer Institute, National Institutes of Health under contract HHSN26120080001E. The content of this publication does not necessarily reflect the views or policies of the Department of Health and Human Services, nor does mention of trade names, commercial products, or organizations imply endorsement by the U.S. Government.

## Author contributions

J. Yoon designed and co-executed experiments with M. Lines with the assistance of K. Murray, A. Luo, T. Yang, and Y. Hwang. J. Yoon, M. Lines, and I. O. Daar wrote the manuscript. I. O. Daar and C. Westlake provided insight on experimental direction. J. Yoon supervised the project. All of the authors discussed the results and commented on the manuscript. This project posthumously honors the work of lab member Kenan Murray.

## Data availability

Imaging data and other data supporting the findings of this study are available from the corresponding author on reasonable request.

---

**Supplementary Figure 1.**
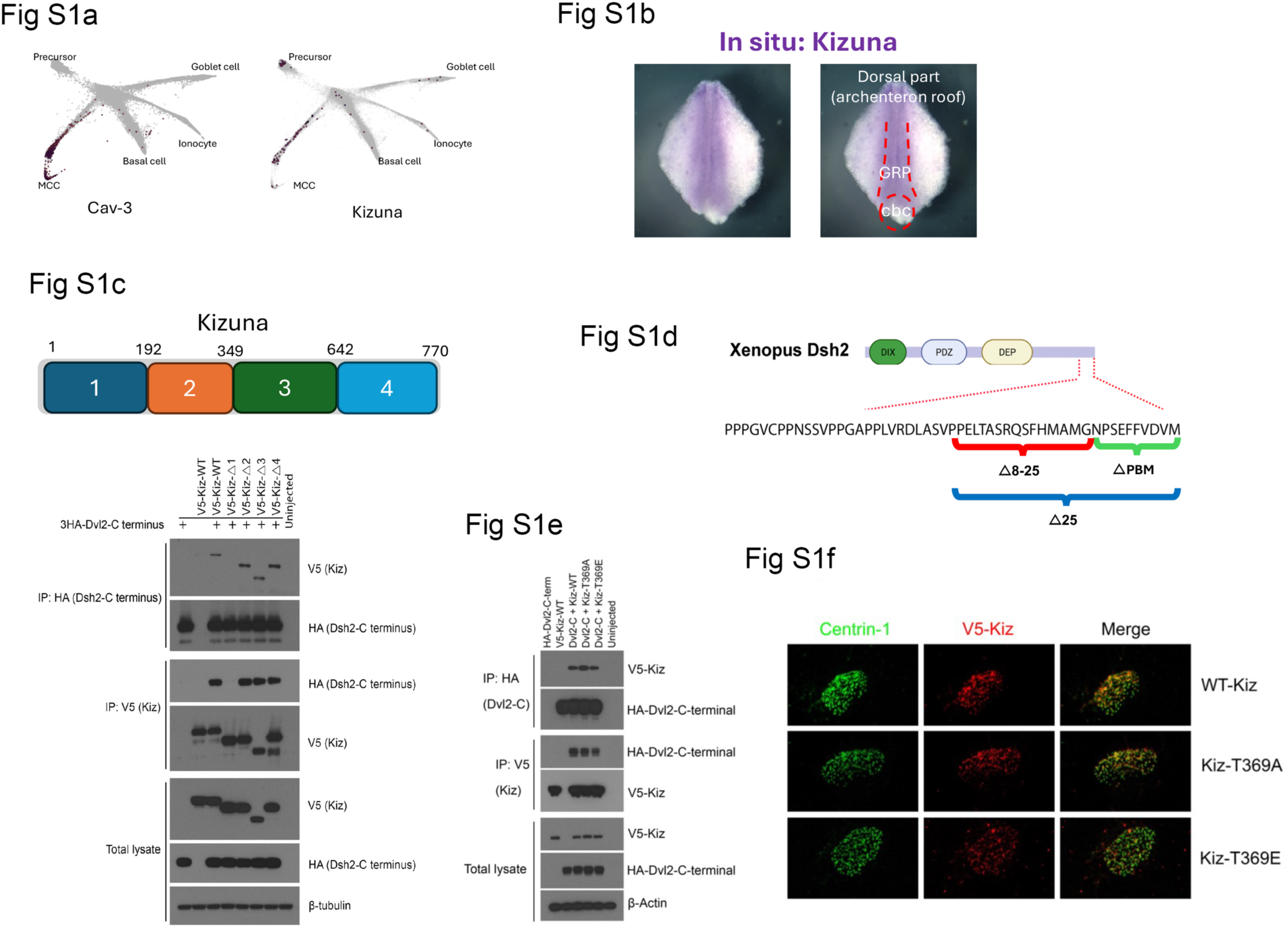
Characterization of Kizuna and its interaction with Dvl. (a) Re-analysis of single-cell RNA-seq data from the Kedar Nath Natarajan laboratory. Cav3 serves as a representative MCC marker. (b) In situ hybridization using anti-Kizuna probe in dissected GRP at stage 20. (c) Schematic representation of Kizuna serial deletion constructs. Co-IP of Dvl-C with Kizuna deletion mutants. (d) Schematic representation of Dvl serial deletion constructs. (e) Co-IP of Dvl-C with Kizuna phosphorylation mutants (T369A or T369E). (f) Co-localization of Centrin and Kizuna phosphorylation mutants (T369A or T369E) in MCCs.

**Supplementary Figure 2.**
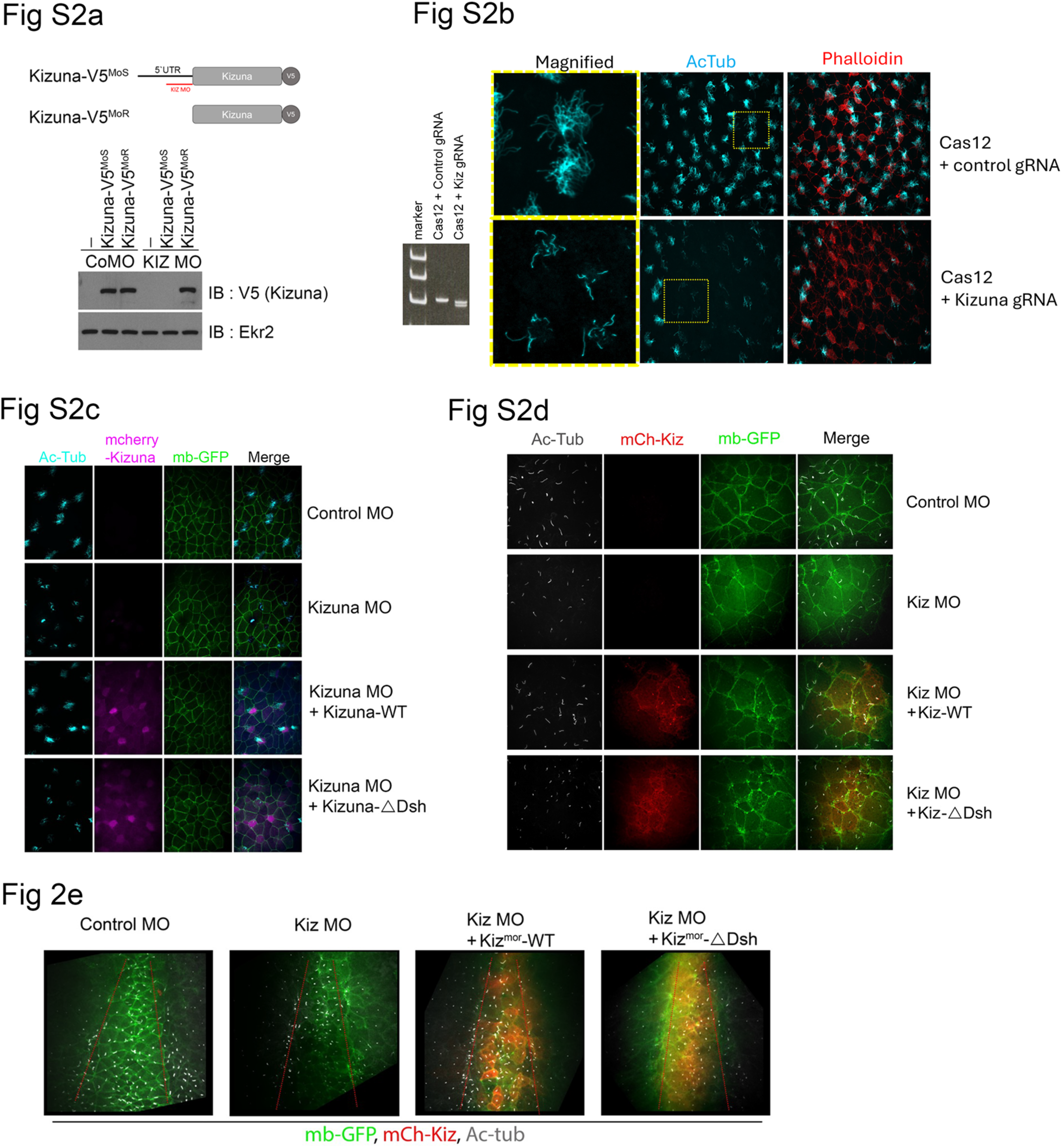
Validation of Kizuna morpholino efficiency and phenotypes. (a) Schematic representation of the Kizuna morpholino target site. Immunoblot confirms MO efficiency in blocking Kizuna expression. (b) Immunostaining of MCCs in Kizuna knockout (KO) embryos at stage 30. (c) Low-magnification images of MCCs in Xenopus embryos at stage 30 following Kizuna MO injection. (d) Immunostaining of mono-motile ciliated cells in the GRP at stage 20. (e) Low-magnification images of mono-motile ciliated cells in the GRP at stage 20.

**Supplementary Figure 3.**
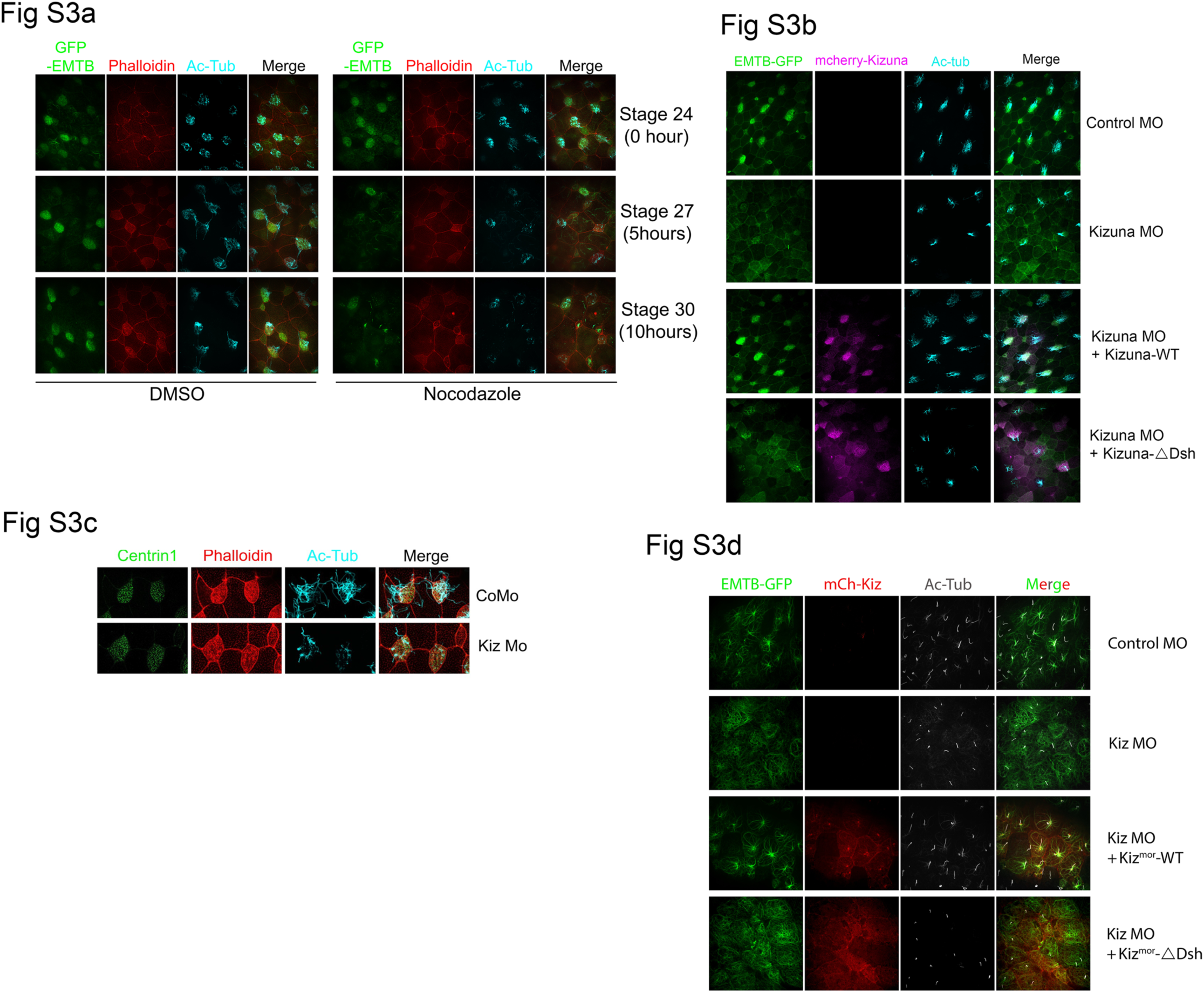
Microtubule organization and actin meshwork in Kizuna morphants. (a) Immunostaining of the apical MT meshwork in MCCs treated with nocodazole (1 μM). (b) Low-magnification images of the apical MT meshwork in MCCs at stage 30 following Kizuna MO injection. (c) Immunostaining of basal bodies and the apical actin meshwork in MCCs at stage 30 following Kizuna MO injection. (d) Low-magnification images of MT nucleation in the GRP at stage 20 following Kizuna MO injection.

**Supplementary Figure 4.**
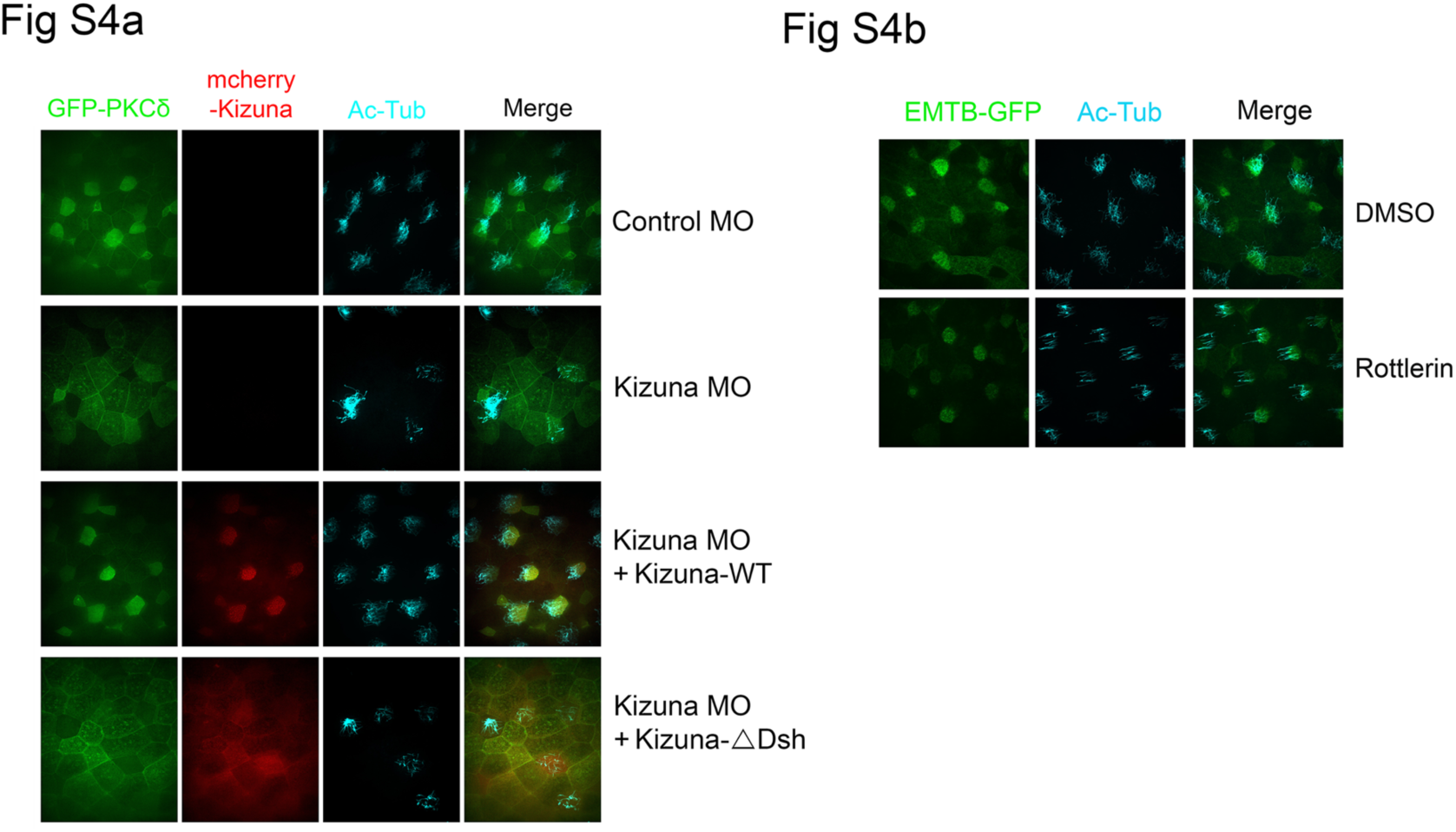
PKCδ localization and rottlerin treatment effects. (a) Low-magnification images showing PKCδ localization in MCCs at stage 30 following Kizuna MO injection. (b) Low-magnification images of the apical MT meshwork in MCCs at stage 30 following rottlerin (10 μM) treatment.

**Supplementary Figure 5.**
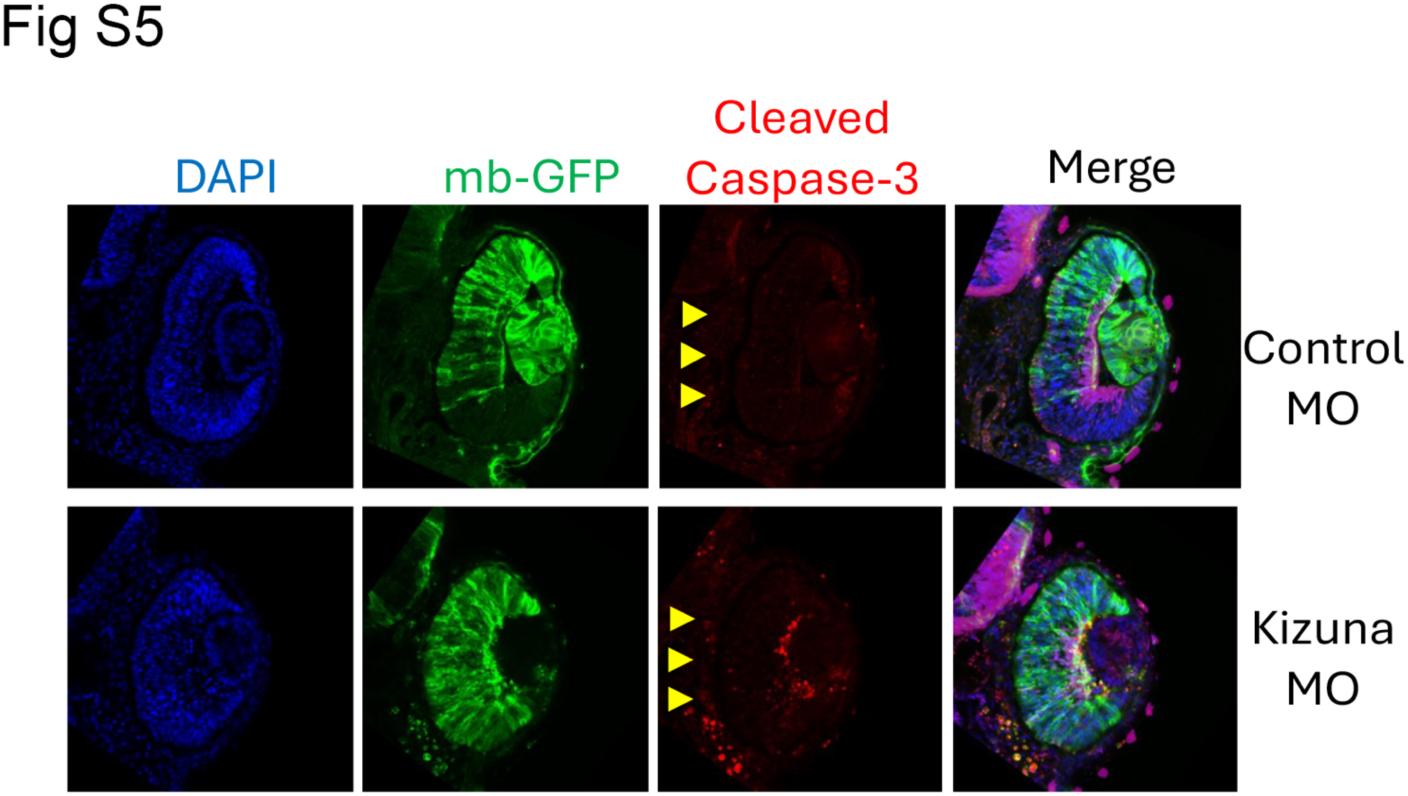
Apoptosis analysis in Kizuna morphants. Immunostaining of the apoptosis marker cleaved caspase-3 in eyes of Xenopus embryos at stage 35 following Kizuna MO injection.

**Supplementary Movie 1.**

Time-lapse imaging of membrane-GFP in MCCs of Xenopus embryos at stage 30. Kizuna MO was injected into the V1.1 blastomere at the 8-cell stage. One frame was captured every 100 ms.

**Supplementary Movie 2.**

Time-lapse imaging of membrane-GFP in mono-motile ciliated cells of the GRP at stage 20. Kizuna MO was injected into the two dorsal marginal zones at the 4-cell stage. One frame was captured every 100 ms.

